# A metagenomic approach to One Health surveillance of antimicrobial resistance in a UK veterinary centre

**DOI:** 10.1101/2025.02.12.636430

**Authors:** Linzy Elton, Alonso Dupuy Mateos, Siân Marie Frosini, Rosanne Jepson, Sylvia Rofael, Timothy D McHugh, Emmanuel Q Wey

## Abstract

**Introduction:** There are currently no standardised guidelines for genomic surveillance of One Health (OH) antimicrobial resistance (AMR). This project aimed to utilise metagenomics to identify AMR genes present in a companion animal hospital and compare these with phenotypic results from bacterial isolates from clinical specimens from the same veterinary hospital.

**Methods:** Samples were collected from sites around a primary companion animal veterinary hospital in North London. Metagenomic DNA was sequenced using Oxford Nanopore Technologies (ONT) MinION. The sequencing data were analysed for AMR genes, plasmids and clinically relevant pathogen species. These data were compared to phenotypic speciation and antibiotic susceptibility tests (ASTs) of bacteria isolates from patients.

**Results:** The most common resistance genes identified were *aph* (n=101 times genes were isolated across 48 metagenomic samples), *sul* (84), *bla*_CARB_ (63), *tet* (58) and *bla*_TEM_ (46). In clinical isolates, a high proportion of phenotypic resistance to the β-lactams was identified. Rooms with the greatest mean number of resistance genes identified per swab site were the medical preparation room, dog ward and surgical preparation room. Twenty-four Gram-positive and four enterobacterial plasmids were identified. Sequencing reads matched with 14/22 (64%) of the phenotypically isolated bacterial species.

**Discussion:** Metagenomics identified AMR genes, plasmids and species of relevance to human and animal medicine. Communal animal-handling areas harboured more AMR genes than areas animals did not frequent. When considering infection prevention and control (IPC) measures, adherence to, and frequency of, cleaning schedules, alongside potentially more comprehensive disinfection of animal-handling areas may reduce the number of potentially harmful bacteria present.

PubMed “veterinary” or “companion” AND “AMR” or “resistan*” NOT (Review[Publication Type]) (2018-2023)

“veterinary” or “companion” AND “AMR” or “resist*” AND “sequencing” or “metagenomic*” “veterinary” or “companion” AND “AMR” or “resistan*” NOT (Review[Publication Type]) (2018-2023)

https://www.iscaid.org/clinical-practice

## Introduction

Globally, many people live alongside companion or smallholding animals. In the UK, half of adults own a pet (1). The concept of One Health (OH) has expanded the focus from human healthcare settings to include animal populations, wastewater and the environment as sources of antimicrobial resistance (AMR) transmission (2–4). AMR was associated with 4.95 million human deaths in 2019, but data on the impact of AMR on companion animal morbidity and mortality is largely missing (5).

Despite the increasing popularity of companion animals, within the veterinary setting AMR surveillance focus has been largely on food production animals. Multi-drug-resistant organisms (MDROs) such as methicillin-resistant staphylococci (MRS), Extended-spectrum beta-lactamases (ESBLs) and Carbapenem-resistant Enterobacteriales (CREs) demonstrate resistance to multiple, important antibiotic groups. MDROs commonly cause human-healthcare associated infections (HHAIs) and are also increasingly being isolated from animals and the environment (6). Companion animals such as dogs and cats are now considered to be AMR reservoirs (7,8).

Studies have shown that important MDROs such as *Escherichia coli*, *Staphylococcus aureus* and *Staphylococcus pseudintermedius* have been isolated from veterinary hospital environments around the world (9–11). A recent systematic review of human-healthcare associated infections (HHAI) and zoonotic MDROs stated that clinically relevant species were reportedly identified from environmental surfaces, clinical infections and fomites such as mobile phones and equipment such as hair clippers (11). *Staphylococcus* species (spp.) were the most reported organisms across the studies, followed by *E. coli*, *Enterococcus* spp., *Salmonella* spp., *Acinetobacter baumannii*, *Clostridium difficile* and *Pseudomonas aeruginosa* (11). Veterinary healthcare workers, like their human healthcare counterparts, have also been found to be colonised with MDROs such as methicillin-resistant *Staphylococcus aureus* (MRSA) (12). Jobs involving animal handling are a risk factor for carriage of MDROs (11,13,14). There is also a greater risk of becoming infected (requiring treatment) with an MDRO if a host has been previously asymptomatically colonised (15). Companion animals can become long-term carriers of MDROs following clinical infection and thus contribute to transmission within the animal, human healthcare and community environment (16–19).

Most of the previous literature looking at MDROs and AMR transmission in clinical veterinary and food chain environments has utilised phenotypic testing to identify both species and AMR. Several clinical studies in human hospitals have utilised genomics to show that AMR determinants and MDRO reservoirs can be found in healthcare setting environments, such as wastewater and from fomite swabbing, with similar data from food-animal production systems (20–24). There is little data to show whether these AMR reservoirs are found in primary companion animal veterinary hospital environments.

Standardising tools and guidelines for the genomic surveillance of One Health AMR is complicated, owing to the varying pathogens, sample types and hosts (25). In the UK, whilst there are human clinical and farm animal AMR surveillance programmes, there is no national surveillance programme for AMR in companion animals, and only limited data exists (26,27). In this project we aimed to utilise a metagenomic approach to identify AMR genes and clinically (both human and animal) relevant species present in a primary companion animal hospital environment and compare and relate them to phenotypic speciation and antibiotic susceptibility test (AST) results.

## Methods

### Clinical data collection

Phenotypic pathogen species and AST data from all patient samples (dog, cats, rabbits) submitted for bacteriology culture and susceptibility testing between January-December 2022 were obtained from the veterinary practice clinical records. Results obtained from the practice’s laboratory information management system (LIMS) included all samples irrespective of source (e.g. urine, bile, skin, wounds, faeces etc.) submitted for bacteriology (IDEXX, Wetherby, UK). Bacterial speciation and minimum inhibitory concentrations (MICs) results, interpreted from CLSI VET01S ED5:2020 guidelines and using VITEK MS MALDI-TOF and VITEK-2 MIC, were extracted from the LIMS linked to IDEXX (28).

### Clinical data analysis

Clinical data were descriptively analysed to display the relationships between clinical isolate species, drug resistances seen and patient species.

### Environmental site selection and sample collection

For environmental sampling, a site visit to the veterinary hospital was undertaken prior to sample collection in order to evaluate room layout (Supplementary Materials S1), staff and patient routes of travel, and determine prioritised sampling locations including both patient and non-patient areas (Supplementary Materials S2). Where there were multiple same-use rooms, e.g. consultation rooms, one representative location was chosen. Non-clinical rooms were also evaluated, including the visitor toilets. Samples were obtained on a normal working week afternoon shift in September 2022 from pre-designated locations. Sample types included wastewater samples (e.g. dog kennel waste drain), sponge swabs of surfaces (e.g. consulting tables), cotton stick swabs (e.g. sink drain holes) and liquid samples (e.g. ultrasound gel).

Further details of swabbing procedures for each swabbing site can be found in Supplementary Materials S3. For wastewater samples, 500 µL of wastewater was collected using a Pasteur pipette from the waste pipes of sinks, then placed into sterile screwcap tubes pre-dosed with 500 µL DNA/RNA Shield (Zymo Research Corporation). Cotton-tipped stick swabs (SS352, Appleton Woods) pre-moistened with 1x PBS (J61196.AP, Thermo Scientific) and stored in 500 µL PBS and 500 µL DNA/RNA Shield, and sponge swabs (TS/15-B, Technical Service Consultants Ltd.), pre-dosed with neutraliser buffer (29), were used to collect samples from approximately 5 cm^2^ of hard surfaces. Stick swab samples were transferred to sterile universal tubes containing 500 µL of DNA/RNA Shield. Sponge swabs were stored in their accompanying pouches until processing in the laboratory, where the liquid was aseptically removed with a Pasteur pipette and 500 µL was added to 500 µL DNA/RNA Shield. All samples were refrigerated (2-8°C) within 2 hours of collection and processed in the laboratory within 24 hours.

### DNA extraction and quantification

DNA was extracted from all environmental samples using the ZymoBIOMICS™ DNA Miniprep Kit (Zymo Research Corporation) following manufacturer’s instructions (30). A sample of ZymoBIOMICS™ Microbial Community Standard (Zymo Research Corporation) was included as an extraction control, following manufacturer’s instructions (31). DNA was assessed for concentration using the Qubit™ dsDNA BR Assay Kit (Thermo Fisher).

### Library preparation

DNA libraries were prepared using the ONT Rapid PCR Barcoding Kit (SQK-RPB004) with a DNA input of 1-5 ng and following the manufacturers’ instructions (32). ZymoBIOMICS™ Microbial Community DNA Standards (Zymo Research Corporation) were included as a sequencing control, following manufacturer’s instructions (33).

### Sequencing and basecalling

Libraries were loaded onto a flow cell, version R9.4.1, (Oxford Nanopore Technologies (ONT)) using a MinION device and were sequenced for 72 hours, using the default parameters on the MinKNOW software (v23.04.6). Basecalling was performed by Guppy integrated into the MinKNOW software [41], using the high accuracy algorithm. A total of 48 independent samples were evaluated in 4 runs of 12 barcoded samples. Two samples obtained from ultrasound gel were combined (samples 46 and 47) and the sample obtained from petroleum jelly did not yield any DNA and therefore could not be sequenced.

### Sequencing data analysis

Of the 48 samples sequenced, nine did not harbour any reads. Fastq files were quality checked using FastQC (v0.21.1) and MultiQC (v1.15) (34,35). Barcodes were trimmed from the reads using Guppy (v6.5.7) (36). Sample data were analysed for the presence of AMR genes using KmerResistance 2.2 (v02-2018), with 70% identity threshold (37,38). Plasmids were identified using PlasmidFinder 2.1 (v2.0.1), with a minimum identity threshold of 95% and minimum coverage threshold of 60% (37,39) and for species using Kraken (v2.1.3) (40). Sequence data were deposited in European Nucleotide Archive (ENA) under BioProject PRJEB84924 and outlined in Supplementary Materials S2. For analysis of clinical isolate AMR status, an isolate was deemed multi-drug resistant (MDR) if it showed resistance to antibiotics in three or more antibiotic classes, to which the bacterial species does not show intrinsic resistance (41,42).

## Results

### Clinical data

The AST and speciation results for 148 clinical bacterial isolates from 91 dogs, 56 cats and 1 rabbit were obtained (Table 1). Twenty-two bacterial species or genera were identified, of which the most commonly isolated were *Escherichia coli* (46 isolates), *Staphylococcus pseudintermedius* (28 isolates), *Pseudomonas aeruginosa* (13 isolates), *Enterococcus faecalis* (12 isolates) and *Pasteurella multocida* (10 isolates). From AST data, the mean frequency of resistance calls per isolate across all species was 2.6 (SD=33.5). Species with the highest number of mean resistances were *Staphylococcus saprophyticus* (12, SD=0.0), *Staphylococcus haemolyticus* (10, SD=0.0), *S. aureus* (9, SD=5.0), *Staphylococcus lentus* (now *Mammaliicoccus lentus*) (43) (9, SD=0.0) and *Morganella morganii* (7, SD=0.0). Those with the lowest mean resistances were *P. multocida*, *P. aeruginosa*, Salmonella spp. and *Serratia fonticola* (0, SD=0.0). *P. aeruginosa* had the highest number of mean intermediate resistances (1.2, SD=0.4).

**Table 1.**
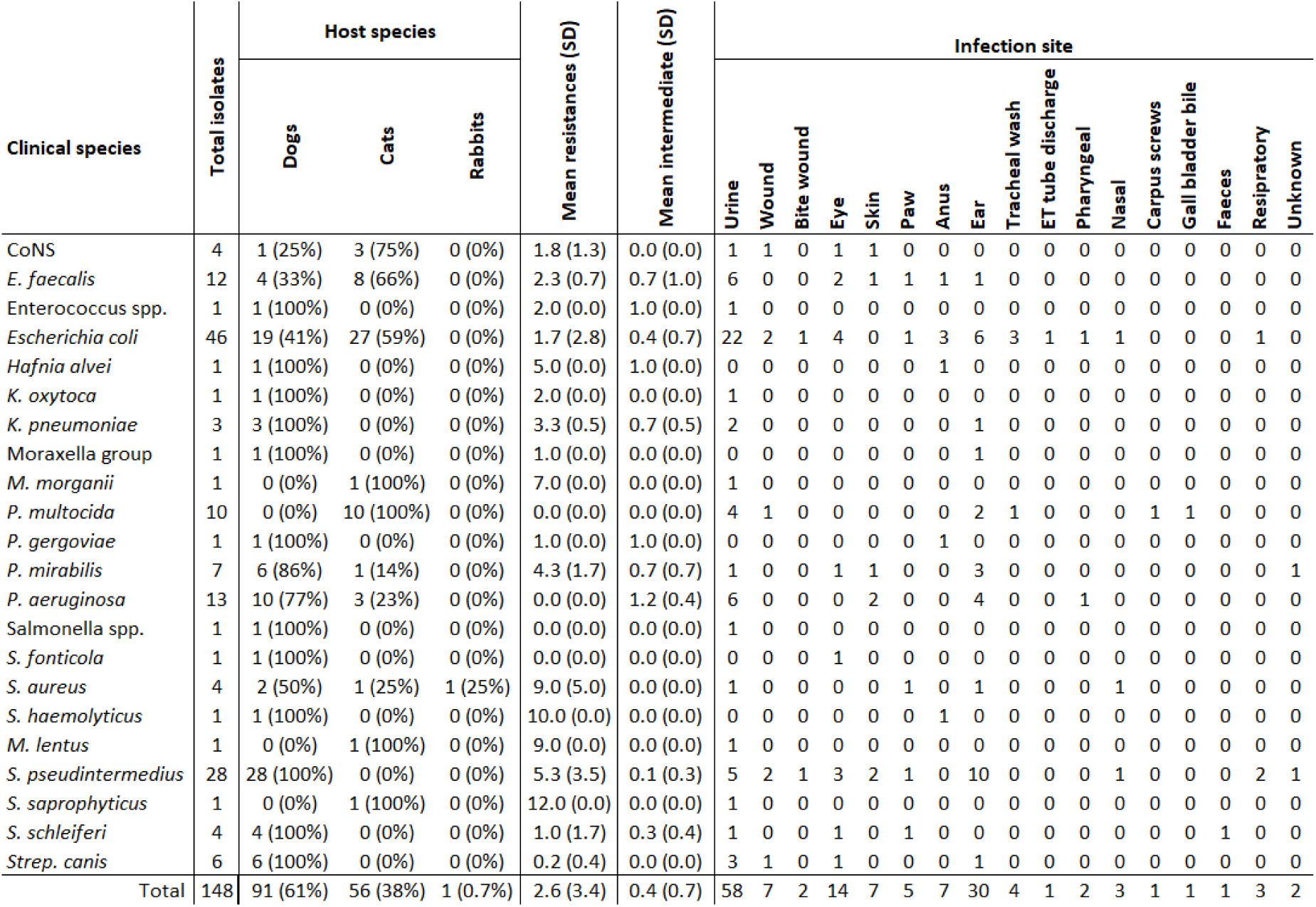
Bacterial isolates identified from clinical samples during the year 2022. Total number of isolates identified, number of isolates stratified by host species, mean number of resistances per isolate and mean number of intermediate resistances per isolate. CoNS = Coagulase negative Staphylococci, SD = standard deviation, ET = endotracheal.

The clinical isolates were most commonly found to have resistance to drugs in the β-lactam group; 35/148, (23.8%) were resistant to at least one β-lactam antibiotic (Table 2). The drugs with the highest percentages of isolate resistance were penicillin with 33/61 isolates tested (54.1%), amoxicillin 57/119 isolates tested (47.9%), ampicillin 56/126 (44.4%), cephalexin 31/117 (26.5%) and cefovecin 27/128 (21.1%). Resistance was not identified, when tested to, florfenicol (0/56), moxifloxacin (0/6), mupirocin (0/39) or rifampicin (0/39).

**Table 2.** Phenotypic AST results from 148 bacterial isolates by VITEK minimum inhibitory concentration testing from patient samples. ICR = inducible clindamycin resistance.

| Drug group | Antibiotic | Total tested | Resistant | Intermediate |
| --- | --- | --- | --- | --- |
| Aminoglycoside | Amikacin | 118 | 9 (7.6%) | 0 (0.0%) |
|  | Gentamicin | 118 | 2 (1.7%) | 2 (1.7%) |
|  | Neomycin | 110 | 5 (4.5%) | 2 (1.8%) |
| β-lactams | Penicillin | 61 | 33 (54.1%) | 0 (0.0%) |
|  | Ampicillin | 126 | 56 (44.4%) | 0 (0.0%) |
|  | Amoxicillin | 119 | 57 (47.9%) | 0 (0.0%) |
|  | Cloxacillin | 43 | 9 (20.9%) | 0 (0.0%) |
|  | Oxacillin | 43 | 9 (20.9%) | 0 (0.0%) |
|  | Amoxicillin-clavulanic acid | 129 | 18 (14.0%) | 7 (5.4%) |
|  | Cephalexin | 117 | 31 (26.5%) | 0 (0.0%) |
|  | Cefovecin | 128 | 27 (21.1%) | 2 (1.6%) |
|  | Cefpodoxime | 64 | 7 (10.9%) | 0 (0.0%) |
|  | ESBL screen | 50 | 1 (2.0%) | 0 (0.0%) |
| Amphenicols | Chloramphenicol | 125 | 9 (7.2%) | 10 (8.0%) |
|  | Florfenicol | 56 | 0 (0.0%) | 0 (0.0%) |
| Fluoroquinolones | Ciprofloxacin | 141 | 5 (3.5%) | 5 (3.5%) |
|  | Enrofloxacin | 142 | 5 (3.5%) | 8 (5.6%) |
|  | Marbofloxacin | 142 | 7 (4.9%) | 2 (1.4%) |
|  | Moxifloxacin | 6 | 0 (0.0%) | 0 (0.0%) |
|  | Pradofloxacin | 115 | 4 (3.5%) | 2 (1.7%) |
| Fusidic acid | Fusidic acid | 39 | 5 (12.8%) | 0 (0.0%) |
| Macrolides | Azithromycin | 49 | 10 (20.4%) | 0 (0.0%) |
|  | Erythromycin | 49 | 10 (20.4%) | 0 (0.0%) |
| Lincosamides | Clindamycin | 24 | 4 (16.7%) | 0 (0.0%) |
|  | ICR test | 16 | 3 (18.8%) | 0 (0.0%) |
|  | Lincomycin | 39 | 9 (23.1%) | 0 (0.0%) |
| Sulphonamides | Potentiated sulphonamides | 122 | 15 (12.3%) | 0 (0.0%) |
| Tetracyclines | Tetracycline | 79 | 16 (20.3%) | 0 (0.0%) |
|  | Doxycycline | 85 | 19 (22.4%) | 0 (0.0%) |
|  | Minocycline | 56 | 3 (5.4%) | 0 (0.0%) |
| Mupirocin | Mupirocin | 39 | 0 (0.0%) | 0 (0.0%) |
| Nitrofurantoin | Nitrofurantoin | 78 | 8 (10.3%) | 3 (3.8%) |
| Polypeptide | Polymyxin B | 13 | 0 (0.0%) | 13 (100.0%) |
| Rifampicin | Rifampicin | 39 | 0 (0.0%) | 0 (0.0%) |

One of the 46 *E. coli* (2%) isolates obtained from a feline ear swab was identified as MDR by phenotypic AST and 1/7 (14%) *P. mirabilis* isolate from a canine ear swab. Two of the four *S. aureus* isolates (50%) were classified as MRSA, identified from a canine nasal swab and feline ventral wound. Methicillin resistant *Staphylococcus pseudintermedius* (MRSP) was identified in 4/28 (14%) *S. pseudintermedius* isolates (three canine ear swabs and one canine wound swab) and a further 6/28 (21%) were classed as MDR (one canine ear swabs, two canine eye swabs, two canine skin swabs and one canine urine sample), albeit methicillin susceptible. The single *S. saprophyticus*, from a feline urine sample, was classed as MDR. Single isolates of *M. lentus* (feline ear swab) and *Staphylococcus haemolyticus* (canine carpel screws) were also considered as MDR. A breakdown of clinical AST results stratified by species can be found in Table 3. A list of innate or commonly described resistances for the clinical species analysed in this study can be found in Supplementary Materials S4.

**Table 3.** Phenotypic resistances from clinical isolates. I = intermediate resistance, NT = not tested, ICR = inducible clindamycin resistance. Blue indicates an antibiotic was not tested for that bacterial species, white indicates 0% resistance, pink to red indicates increasing numbers of isolates with resistance to each antibiotic. Orange indicates the number of isolates with only intermediate resistance to each antibiotic. Yellow indicates the number of isolates with resistance and also intermediate resistance.

| Species | Total isolates | Aminoglycosides |  |  |  | β lactams |  |  |  |  |  |  |  |  | Amphenicols |  | Fluoroquinolones |  |  |  |  |
| --- | --- | --- | --- | --- | --- | --- | --- | --- | --- | --- | --- | --- | --- | --- | --- | --- | --- | --- | --- | --- | --- |
|  |  | Gentamicin | Neomycin | Amikacin | Azithromycin | Ampicillin | Amoxicillin-clavulanic acid | Amoxicillin | Cloxacillin | Oxacillin | Penicillin | Cefovecin | Cephalexin | Cefpodoxime | Chloramphenicol | Florfenicol | Ciprofloxacin | Enrofloxacin | Marbofloxacin | Moxifloxacin | Pradofloxacin |
| Coagulase neg. Staph. | 4 | 0% | 0% | 100% | 0% | 25% | 0% | 25% | 0% | 0% | 25% | 0% | 0% | NT | 0% | 0% | 0% | 0% | 0% | NT | 0% |
| *E. faecalis* | 12 | NT | NT | NT | NT | 0% | 0% | 0% | NT | NT | 0% | 83% | 100% | NT | 0% | 0% | 25% (I) | 25% (I) | 17% (17% I) | NT | NT |
| Enterococcus spp | 1 | NT | NT | NT | NT | NT | 0% | NT | NT | NT | 0% | 100% | 100% | NT | 0% | 0% | 0% | 0% | 0% | NT | NT |
| *E. coli* | 46 | 0% | 2% | 0% | NT | 33% | 17% (17% I) | 33% | NT | NT | NT | 13% | 9% | 9% | 4% (17% I) | NT | 4% (4% I) | 4% (4% I) | 4% | NT | 4% (2% I) |
| *H. alvei* | 1 | 0% | 0% | 0% | NT | 100% | 100% | 100% | NT | NT | NT | 100% (I) | 100% | 100% | 0% | NT | 0% | 0% | 0% | NT | 0% |
| *K. oxytoca* | 1 | 0% | 0% | 0% | NT | 100% | 0% | 100% | NT | NT | NT | 0% | 0% | 0% | 0% | NT | 0% | 0% | 0% | NT | 0% |
| *K. pneumoniae* | 3 | 0% | 0% | 0% | NT | 100% | 100% | 100% | NT | NT | NT | 0% | 33% | 0% | 0% | NT | 0% | 0% | 0% | NT | 0% |
| Moraxella group | 1 | 0% | NT | 0% | NT | 0% | 0% | NT | NT | NT | NT | 100% | NT | NT | 0% | NT | 0% | 0% | 0% | NT | NT |
| *M. morganii* | 1 | 0% | 0% | 0% | NT | 100% | 100% | 100% | NT | NT | NT | NT | 100% | NT | 0% | NT | 0% | 0% | 0% | NT | 0% |
| *P. multocida* | 10 | NT | 0% | NT | NT | 0% | 0% | 0% | NT | NT | NT | 0% | NT | NT | NT | NT | 0% | 0% | 0% | NT | 0% |
| *P. gergoviae* | 1 | 0% | 0% | 0% | NT | NT | 0% | NT | NT | NT | NT | 0% | 100% | 0% | 100% (I) | NT | 0% | 0% | 0% | NT | 0% |
| *P. mirabilis* | 7 | 0% | 0% | 0% | NT | 43% | 100% (I) | 43% | NT | NT | NT | 14% (I) | 14% | 0% | 14% (14% I) | NT | 0% | 0% | 0% | NT | 0% |
| *P. aeruginosa* | 13 | 0% | NT | 0% | NT | NT | NT | NT | NT | NT | NT | NT | NT | NT | NT | NT | 0% | 23% (I) | 0% | NT | NT |
| Salmonella spp | 1 | NT | 0% | NT | NT | 0% | 0% | 0% | NT | NT | NT | 0% | NT | 0% | 0% | NT | 0% | 0% | 0% | NT | 0% |
| *S. fonticola* | 1 | 0% | 0% | 0% | NT | NT | 0% | NT | NT | NT | NT | 0% | 0% | NT | 0% | NT | NT | 0% | 0% | NT | 0% |
| *S. aureus* | 4 | 50% | 0% | 100% | 25% | 100% | 50% | 100% | 50% | 50% | 100% | 50% | 50% | NT | 25% | 0% | 0% | 0% | 0% | NT | 25% |
| *S. haemolyticus* | 1 | 0% | 0% | 100% | 0% | 100% | 100% | 100% | 100% | 100% | 100% | 100% | 0% | NT | 100% | 0% | 0% | 0% | 0% | NT | 0% |
| *S. lentus* | 1 | 0% | NT | 0% | 0% | 100% | 100% | 100% | 100% | 100% | 100% | 100% | 100% | 100% | 0% | 0% | 0% | 0% | 0% | NT | 0% |
| *S. pseudintermedius* | 28 | 7% (I) | 14% (2% I) | 0% | 29% | 79% | 14% | 82% | 14% | 14% | 82% | 14% | 14% | 0% | 18% | 0% | 7% | 7% | 7% | NT | 7% |
| *S. saprophyticus* | 1 | 0% | NT | 0% | 100% | 100% | 100% | 100% | 100% | 100% | 100% | 100% | 100% | 100% | 0% | 0% | 0% | 0% | 0% | NT | 0% |
| *S. schleiferi* | 4 | 0% | 0% | 0% | 0% | 0% | 0% | 0% | 0% | 0% | 0% | 0% | 0% | NT | 0% | 0% | 25% | 25% | 25% | NT | 25% (I) |
| *S. canis* | 6 | NT | NT | NT | 0% | 0% | NT | NT | NT | NT | 0% | NT | NT | NT | 0% | NT | NT | NT | NT | 0% | NT |

| Species | Total isolates | Fusidic acid | Lincosamides |  | Macrolide | Sulphonamides | Tetracyclines |  |  | Mupirocin | Nitrofurantoin | Polypeptides | Rifamycins | ESBL screen | ICR test |
| --- | --- | --- | --- | --- | --- | --- | --- | --- | --- | --- | --- | --- | --- | --- | --- |
|  |  | Fusidic acid | Clindamycin | Lincomycin | Erythromycin | Potentiated sulphonamides | Doxycycline | Minocycline | Tetracycline | Mupirocin | Nitrofurantoin | Polymyxin B | Rifampicin | ESBL screen | ICR test |
| Coagulase neg. Staph. | 4 | 0% | 0% | 0% | 0% | 0% | NT | 0% | NT | 0% | NT | NT | 0% | NT | NT |
| *E. faecalis* | 12 | NT | NT | NT | NT | NT | 17% | 17% | NT | NT | 0% | NT | NT | NT | NT |
| Enterococcus spp | 1 | NT | NT | NT | NT | NT | 0% | 0% | NT | NT | 100% (I) | NT | NT | NT | NT |
| *E. coli* | 46 | NT | NT | NT | NT | 13% | 15% | NT | 15% | NT | 0% | NT | NT | 2% | NT |
| *H. alvei* | 1 | NT | NT | NT | NT | 0% | 0% | NT | 0% | NT | 0% | NT | NT | NT | NT |
| *K. oxytoca* | 1 | NT | NT | NT | NT | 0% | 0% | NT | 0% | NT | 0% | NT | NT | 0% | NT |
| *K. pneumoniae* | 3 | NT | NT | NT | NT | 0% | 0% | NT | 0% | NT | 66% (I) | NT | NT | 0% | NT |
| Moraxella group | 1 | NT | NT | NT | NT | 0% | 0% | NT | 0% | NT | NT | NT | NT | NT | NT |
| *M. morgani* | 1 | NT | NT | NT | NT | 0% | 100% | NT | 100% | NT | 100% | NT | NT | NT | NT |
| *P. multocida* | 10 | NT | NT | NT | NT | 0% | NT | NT | 0% | NT | NT | NT | NT | NT | NT |
| *P. gergoviae* | 1 | NT | NT | NT | NT | 0% | 0% | NT | 0% | NT | 0% | NT | NT | NT | NT |
| *P. mirabilis* | 7 | NT | NT | NT | NT | 14% | 100% | NT | 100% | NT | 100% | NT | NT | NT | NT |
| *P. aeruginosa* | 13 | NT | NT | NT | NT | NT | NT | NT | NT | NT | NT | 100% (I) | NT | NT | NT |
| Salmonella spp | 1 | NT | NT | NT | NT | 0% | 0% | NT | 0% | NT | NT | NT | NT | NT | NT |
| *S. fonticola* | 1 | NT | NT | NT | NT | 0% | 0% | NT | 0% | NT | 0% | NT | NT | NT | NT |
| *S. aureus* | 4 | 0% | 25% | 0% | 25% | 50% | NT | 25% | NT | 0% | NT | NT | 0% | NT | 25% |
| *S. haemolyticus* | 1 | 100% | 0% | 0% | 0% | 0% | NT | 0% | NT | 0% | NT | NT | 0% | NT | 0% |
| *S. lentus* | 1 | NT | NT | NT | 0% | 0% | 0% | 0% | NT | NT | 0% | NT | NT | NT | NT |
| *S. pseudintermedius* | 28 | 12% | 33% | 29% | 29% | 21% | 50% | 0% | NT | 0% | 0% | NT | 0% | NT | 22% |
| *S. saprophyticus* | 1 | NT | NT | NT | 100% | 0% | 100% | 0% | NT | NT | 0% | NT | NT | NT | NT |
| *S. schleiferi* | 4 | 25% | 0% | 0% | 0% | 0% | NT | 0% | NT | 0% | NT | NT | 0% | NT | 0% |
| *S. canis* | 6 | NT | 0% | NT | 0% | 0% | 0% | NT | 17% | NT | NT | NT | NT | NT | NT |

### Metagenomic data General statistics

Fifty samples were collected from the veterinary hospital environment (34 sponge swabs, seven cotton stick swabs, five wastewater and four liquid samples). Forty-eight samples were sequenced, nine did not harbour any reads. A total of 19 different swabbing sample types across 15 different hospital rooms were evaluated by metagenomic analysis. When the total number of sequencing reads was calculated per sample and stratified by location, the pharmacy had the highest mean number of reads (5,589,075, SD=7,648,964), followed by the medical preparation area (3,299,602, SD=4,727,990) and the surgical preparation area (3,049,427, SD=4,356,555). When stratified by sample type, bin swabs had the highest mean number of reads (3,931,918, SD=5,208,454), followed by consulting tables (3,491,689, SD=4,832,317) and sink plug holes (2,952,655, SD=4,290,089) (see Figure 1). There was no significant difference in the total number of sequencing reads when One-Way ANOVA and Tukey’s multiple comparisons tests were applied to the location, and sample type data.

**Figure 1.**
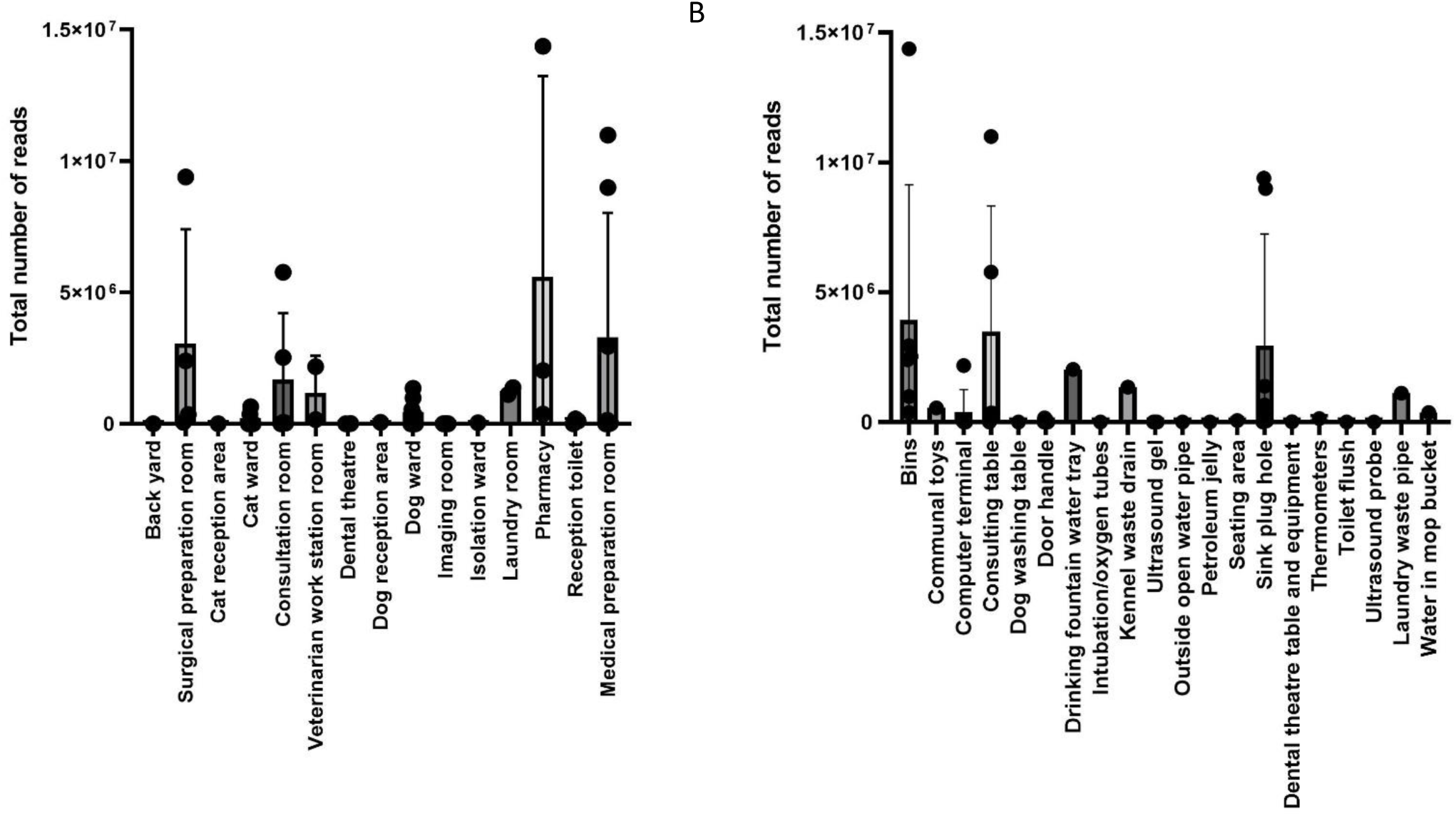
Total number of sequencing reads obtained from metagenomic sequencing stratified by A) location and B) sample type.

Nine of the 48 (18.7%) metagenomic samples did not yield any sequencing reads, so were not included in the proportion analysis (Supplemental Table S2). A further three did not obtain classified reads when analysed and were excluded from further analysis.

When the total number of reads were classified taxonomically into bacterial, human, other eukaryotic, fungal and viral reads, 33/36 (91.6%) samples had >50% bacterial reads (see Figure 2). The three sample sites that did not yield classified reads were the medical preparation area dog wash table (49.0% bacterial, 32.8% human, 16.7% fungal, 1.0% other eukaryote, <0.1% archaea and 0.6% viral), the medical preparation area intubation tubes (10.0% bacterial, 90.3% human, 0.2% fungal, 0.2% other eukaryote and <0.1% archaeal and viral) and medical preparation area thermometers (48.1% bacterial, 51.0% human, 0.1% fungal, 0.1% other eukaryote and <0.1% archaeal and viral). The sample site with the highest proportion of reads identified as fungi was the medical preparation area dog wash table (16.7% of reads). The sample site with the highest proportion of reads identified as viral was the surgical preparation area sink plug hole (36.3% of reads).

**Figure 2.**
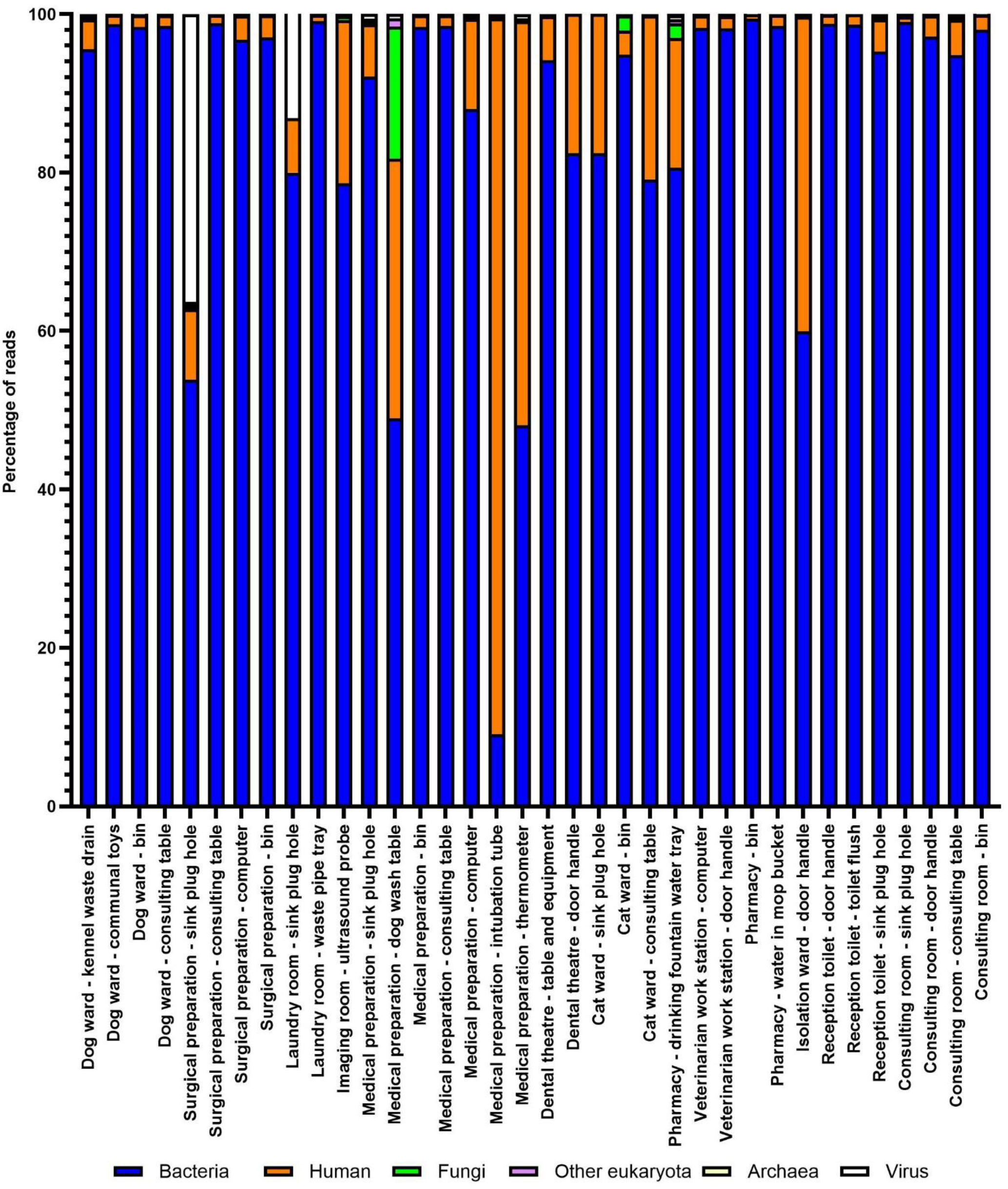
Breakdown of taxonomic classification of metagenomic sample reads. Sample sites that resulted in no reads were not included in this figure.

### AMR gene identification

The most commonly identified AMR genes in the metagenomic samples were *aph*, conferring aminoglycoside resistance (genes were identified 101 times across all metagenomic samples), *sul* conferring sulphonamide resistance (84 genes), *bla*_CARB_ conferring β-lactam resistance (63 genes), *tet* conferring tetracycline resistance (57 genes) and *bla*_TEM_, also β-lactam resistance (46 genes). Ten *mecA* genes were identified, 12 *blaZ* genes, 18 *blaOXA* genes. No *mcr* genes for colistin resistance were identified.

The rooms within the veterinary hospital in which the greatest number of AMR genes identified were the medical preparation area (165 genes, SD=7.3), dog ward (136 genes, SD=6.6), surgical preparation area (63 genes, SD=3.3), veterinarian workstation room (50 genes, SD=2.3) and the consultation room (44 genes, SD=2.2) (Table 4). When stratified by sample type the kennel waste drain (1 sample, 64 genes) showed the greatest number of AMR genes. The consulting tables had the second highest mean (5 samples, mean=29, SD=6.2), then bins (6 samples, mean=20, SD=6.6), sink plug holes (7 samples, mean=16, SD=5.6) and seating areas (2 samples, mean=11, SD=1.9; Table 5).

**Table 4.**
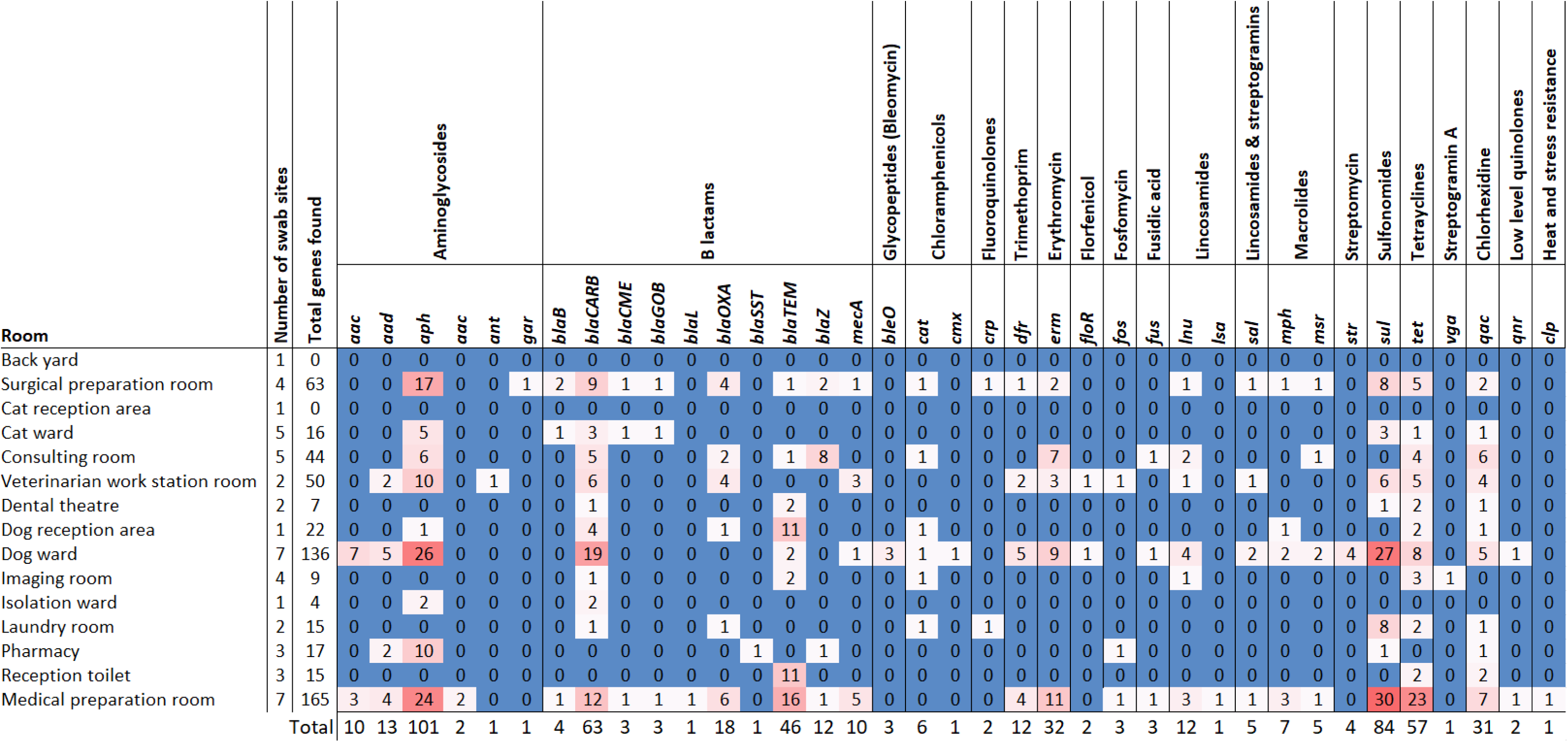
AMR genes stratified by room. Blue indicates that no AMR genes were found, the darker the red, the more genes were identified.

**Table 5.**
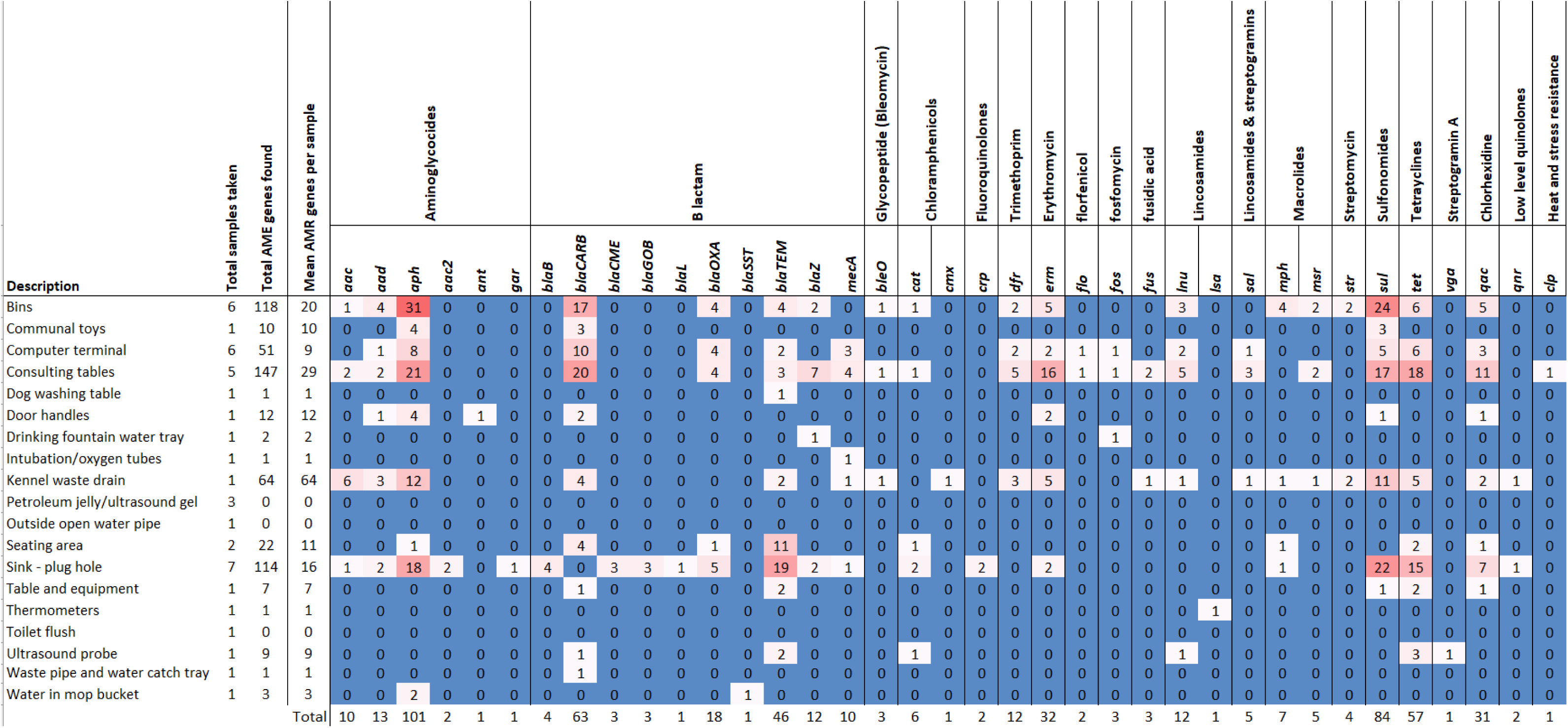
AMR genes stratified by sample type. Blue indicates that no AMR genes were found, the darker the red, the more genes were identified.

### Plasmid identification

Twenty-four plasmids associated with AMR gene transmission in Gram-positives were identified. The highest mean number of Gram-positive plasmids was found in the dog ward (mean=3.7), surgical preparation area (mean=1), consulting room (mean=0.8), medical preparation area (mean=0.6) and the cat ward (mean=0.2) (see Table 6).

**Table 6.**
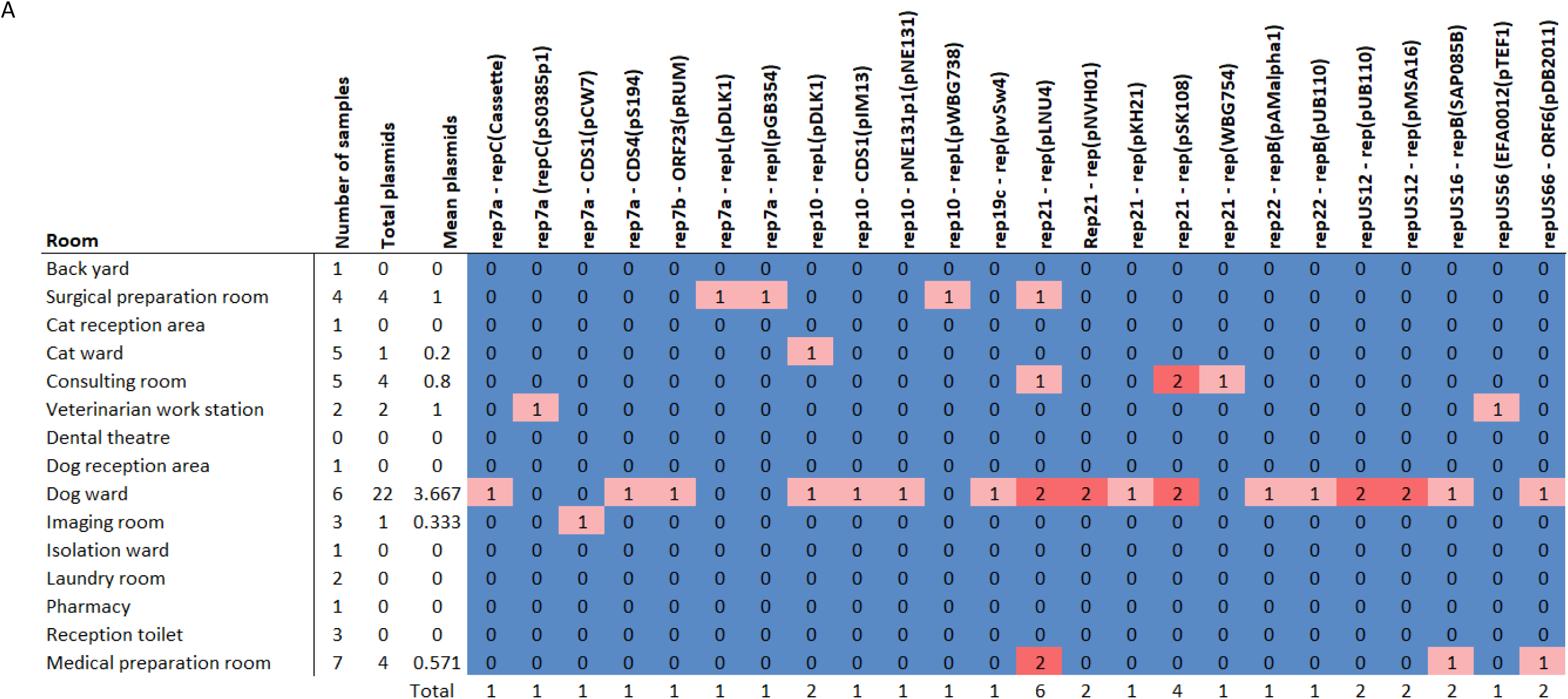
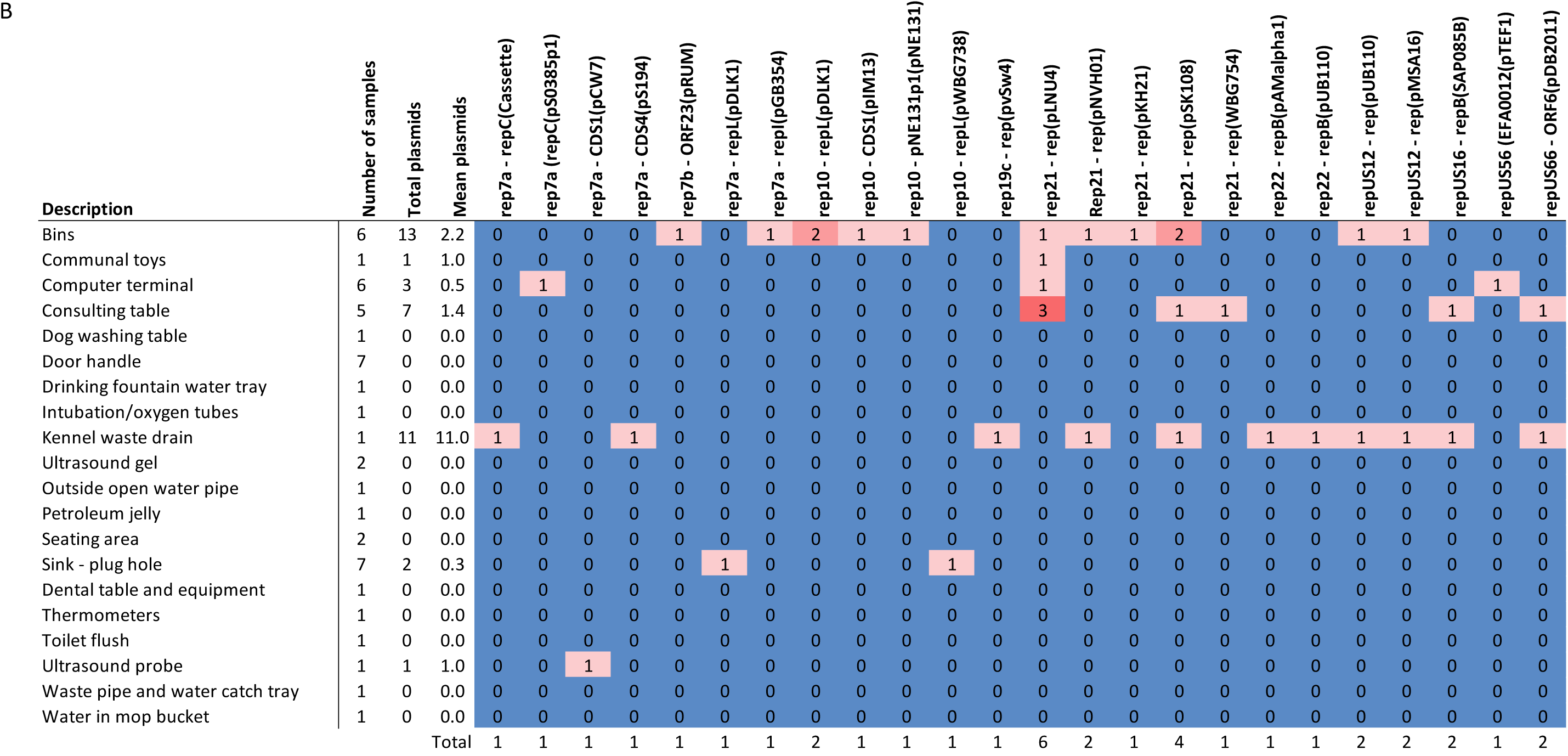
Gram positive plasmids stratified by a) room and b) sample type. Blue indicates that no plasmids were found, the darker the red, the more genes were identified.

Four plasmids associated with AMR gene transmission in Enterobacteriales were identified. Col (pHAD28) was identified from the kennel waste drain swab in the dog ward and Col440I, IncQ2 and IncP6 were identified from the sink plug hole swab from the medical preparation area (see Table 7). No other plasmids associated with Enterobacteriales were identified.

**Table 7.**
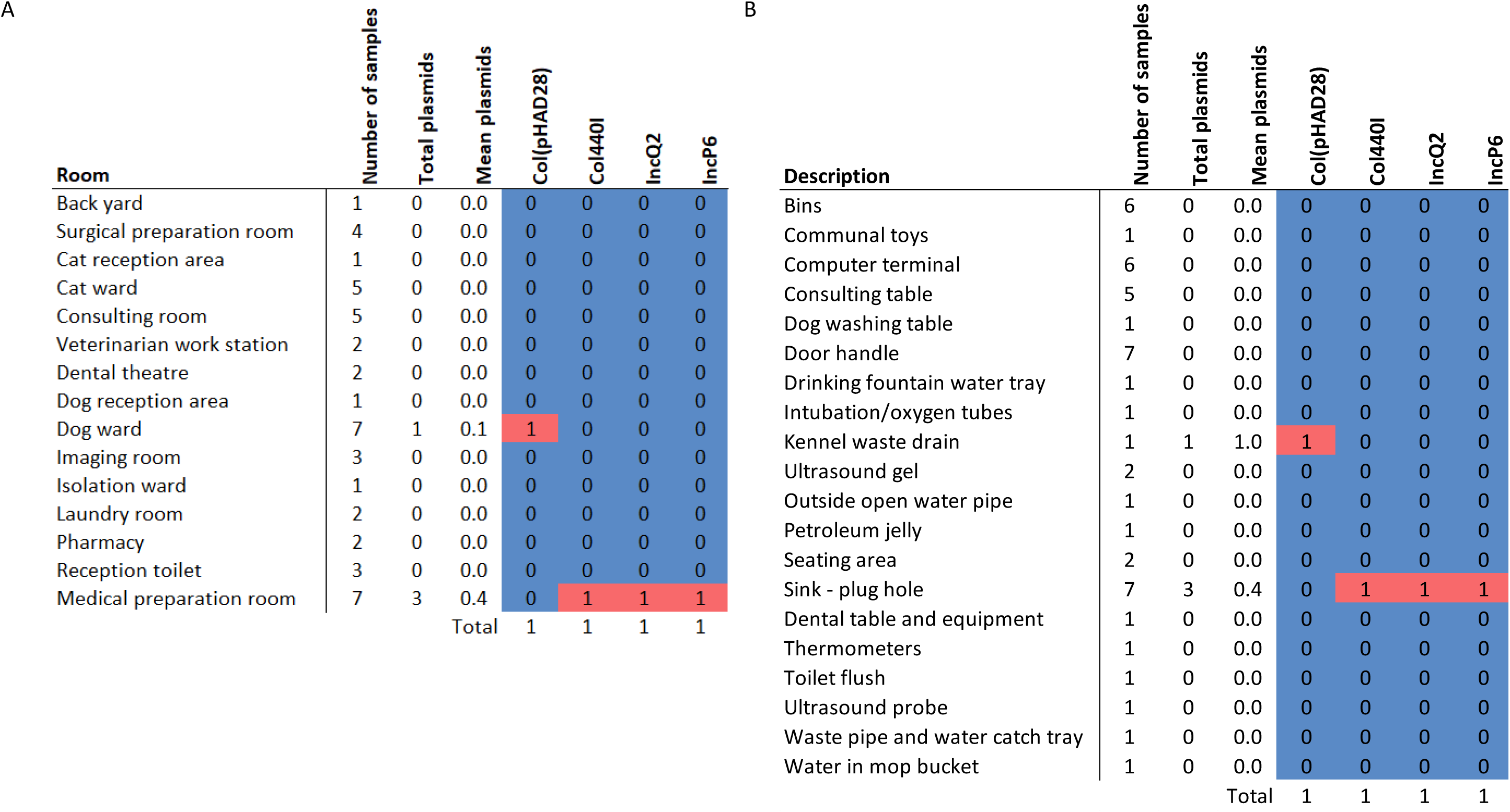
Enterobacterial plasmids stratified by a) room and b) sample type. Blue indicates that no plasmids were found, red indicates that a plasmid was identified.

### Metagenomic speciation

Sequencing reads mapped with 14/22 (64%) of the pathogenic species identified phenotypically from the clinical isolates. The species to which the highest overall mean number of reads mapped was *P. aeruginosa* (1,678,521 reads), *E. coli* (569, 637 reads), the CoNS, for which reads mapped to *S. capitis, S. cohnii, S. epidermidis, S. equorum, S. felis, S. haemolyticus, S. hominis, S. saccharolyticus, S. saprophyticus, Mammaliicoccus sciuri (formerly Staphylococccus sciuri), S. succinus, S. warneri,* and *S. xylosus* (547,367 reads) and the Moraxella group, for which reads mapped to *M. bovoculi, M. catarrhalis, M. cuniculi, M. nonliquefaciens, M. osloensis,* and *M. ovis* (529,572 reads). No sequencing reads mapped to *Hafnia. alvei, Klebsiella oxytoca, P. multocida, Proteus mirabilis, Serratia fonticola, S. auris, S. lentus* or *S. schleiferi*.

Four species (*E. faecalis, E. coli,* Moraxella group and *S. pseudintermedius*) in the clinically pathogenic samples were found across 7/15 (46.7%) locations sampled. Three species (CoNS, *Klebsiella pneumoniae* and *P. aeruginosa*) were found in 6/15 (40.0%) of locations. The medical preparation area had the highest number of reads mapping to species isolated from clinical samples (1,100,313, 30% of the total reads mapped to clinical species), followed by the laundry room (969,382, 26%) and dog ward (541,367, 15%). The back yard, cat reception area, dental theatre and dog reception area had no reads mapping to the clinical species (Table 8A).

**Table 8.**
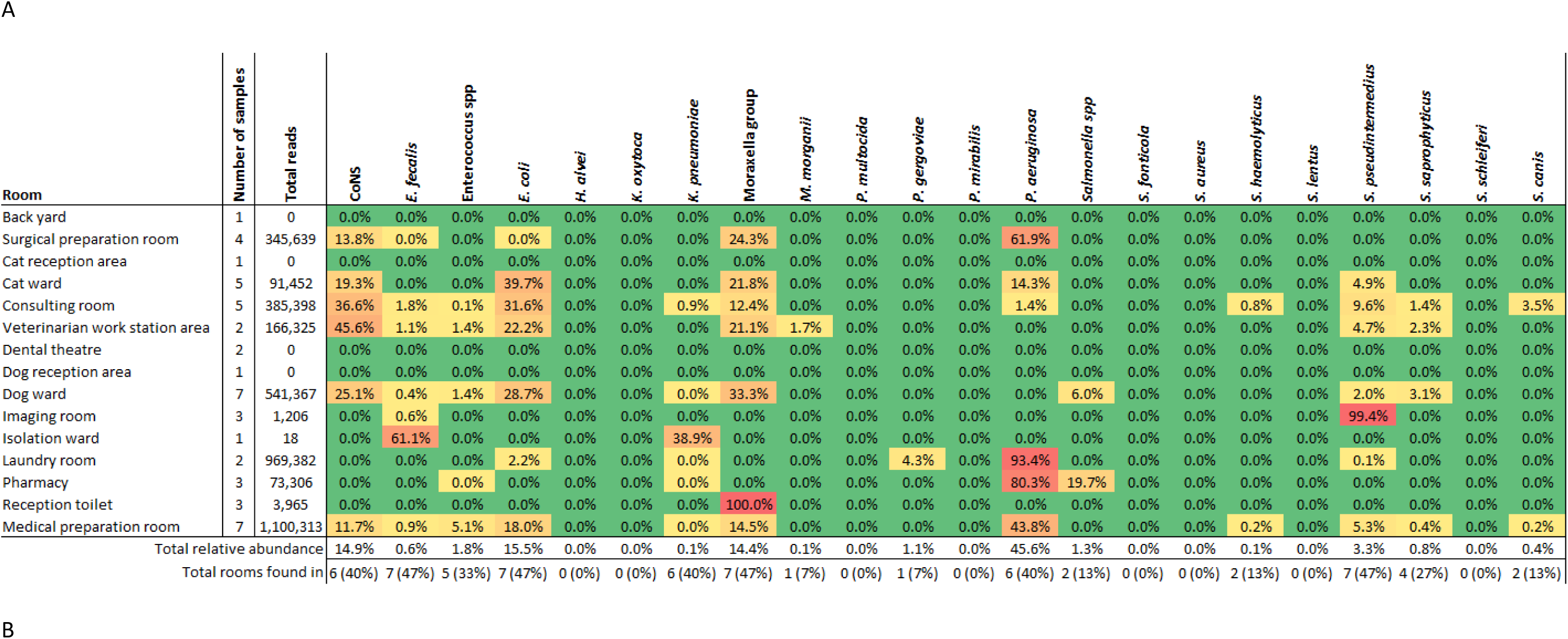
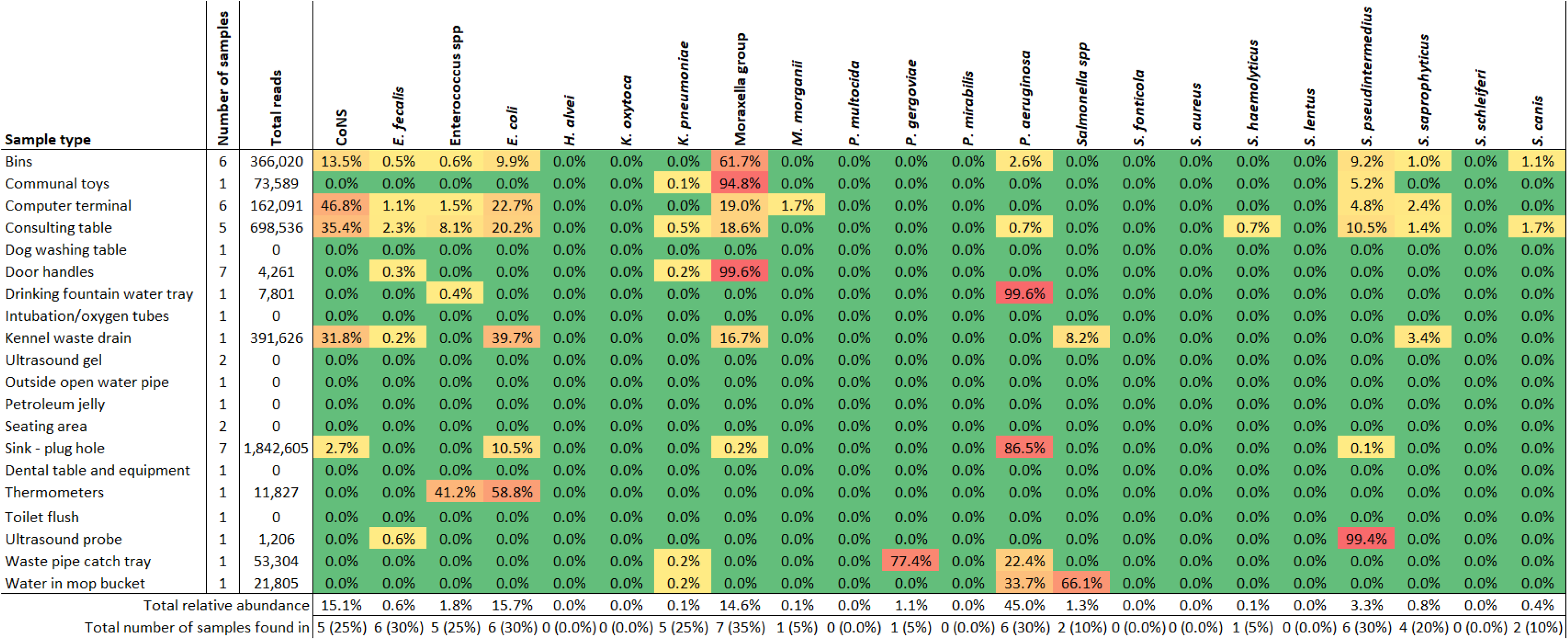
Relative abundance identified for each clinical species. A) relative abundance stratified by location, B) relative abundance stratified by sample type. Species groupings include all species of that genus for which reads were identified. Enterococcus spp (not *E. faecalis*): *E. casseliflavus, E. faecium, E. hirae, E. luminosum*, Moraxella group: *M. bovoculi, M. catarrhalis, M. cuniculi, M. nonliquefaciens, M. osloensis, M. ovis*, Coagulase negative Staphylococci (CoNS): *S. capitis, S. cohnii, S. epidermidis, S. equorum, S. felis, S. haemolyticus, S. hominis, S. saccharolyticus, S. saprophyticus, S. succinus, S. warneri, S. xylosus*, Salmonella spp: *S. enterica.* Green indicates that no AMR genes were found, yellow to red indicates increasing relative abundance.

When stratified by sample type, sink plug hole swabs had the highest number of reads mapping to clinical species (1,842,605, 51% of the total reads mapped to clinical species), followed by the consulting tables (698,536, 19%) and bins (366,020, 10%). The dog washing table, intubation tubes, ultrasound gel, back yard wastewater pipe, petroleum jelly, seating areas, dental surgery table and equipment and toilet flushes had no reads mapping to the clinical species. 97% of reads mapping to *P. aeruginosa* were found in sink plug holes (Table 8B). The Moraxella group was found in 7/20 (35.0%) sample types. *E. faecalis, E. coli P. aeruginosa* and *S. pseudintermedius* were found in 6/20 (30.0%) sample types and CoNS and *K. pneumoniae* were found in 5/20 (25.0%).

## Discussion

In this study we provide evidence that the veterinary hospital environment mirrors human hospitals in that they may act as a reservoir for infectious agents and drug-resistant organisms in particular. AMR genes that were consistent with the phenotypic clinical isolate resistance profiles were found in the environmental samples, and areas with most AMR genes were areas with high levels of animal handling such as the preparation rooms, wards and consultation room, indicating a route to transmission. Bins, consulting tables and sink plug holes were the sample types with the highest numbers of AMR genes identified, also suggesting a link between AMR gene reservoirs and areas with high animal contact and waste disposal. Ten β-lactam resistance genes were identified in the metagenomic samples, and this was echoed in the resistances seen in the clinical isolates. β-lactam antibiotics are clinically important in both animal and human medicine, although access to many of the antimicrobials used to treat ESBL infections in the human sector are not available in veterinary settings (44). Sulphonamide and tetracycline genes were commonly found in the environmental samples (in eight and eleven of the sampling areas respectively), but few clinical isolates showed resistance to these drugs. *mecA* genes were identified in the metagenomic samples and MRSP has been commonly isolated from dogs, including being carried on the skin sub-clinically and long-term following disease resolution, with the potential for dissemination into the environment through natural desquamation (45). MRSP has been increasingly isolated from humans, especially those with close contact with dogs, such as veterinarians and dog owners (46,47).

Plasmids carrying AMR genes are a One Health concern, as they can be easily spread between bacteria, both within and between hosts, and in the environment (48). Most Gram-positive plasmids identified were linked to *Staphylococcus* species and were most commonly recorded in the dog-related areas, such as the dog ward, and from bin samples, suggesting they may be *S. pseudintermedius*-mediated. Many identified contained *erm* genes for erythromycin resistance (including pDLK1, pMSA16, pNE131, pRUM, pWBG738, SAP085B) and nearly a third of the *S. pseudintermedius* phenotypically tested were resistant to erythromycin (49–54). A similar level of resistance to the lincosamides were seen in *S. pseudintermedius*, and pLNU4, encoding *lnu* genes for lincomycin resistance in staphylococci, was the most commonly identified plasmid (55). pSK108 was the second most commonly identified plasmid and along with pWBG574 encodes *qacC*, a multidrug (ß-lactam) resistance gene in staphylococci, which may account for some of the β-lactam resistance seen, alongside *mecA*, which is not plasmid mediated, but found on the chromosomal *SSCmec* cassette (56,57).

Plasmids from the Enterobacteriales were much less commonly identified from the environmental samples and the clinical Enterobacterial isolates generally showed lower rates of phenotypic resistance compared with the Staphylococcal isolates, but all four identified are implicated in the carriage of AMR genes (58–61). Col (pHAD28) plasmids harbour *qnrB* genes, conferring resistance to quinolones, and low numbers of isolates had phenotypic quinolone resistance (62). Col440I can harbour carbapenemase genes such as *bla*_NDM-1_, whilst IncQ2 encodes *bla*_CMY-4_, *bla*_GES-1_ and the sul2-*strA*-*strB* gene cluster (63,64). Whilst resistances to some of the β-lactam genes were common in some of the Enterobacterial isolates, some drugs had low levels of resistance, suggesting a complex interplay of genetic factors. Low numbers of Enterobacterial isolates had sulphonamide resistances and streptomycin was not tested for any of the clinical isolates, likely because streptomycin is not commonly used clinically in companion animals. IncP plasmids have been reported to carry resistance genes for β-lactams, sulphonamides, aminoglycosides and tetracyclines, and recently genes for colistin resistance, although no *mcr* genes were identified in this study (63).

Almost two thirds of the clinically isolated bacterial species were also found in the environmental metagenomic samples, and many are known to also be common environmental species, although pathogenic and non-pathogenic strains may differ phenotypically as a response to differing environmental stressors (65–68). *E. coli* and *P. aeruginosa* were commonly identified as clinical isolates, and also present within the environmental metagenomic samples. *S. pseudintermedius*, the second most frequently found clinical isolate, was commonly found across the locations and sample types in the veterinary hospital. Interestingly, despite it being primarily a canine pathogen, more reads were identified in the cat ward than the dog ward. As it was also found in the preparation rooms where both cats and dogs were handled, this could suggest cross-contamination.

From an infection prevention and control (IPC) perspective, the results identified in this study point to some potential risk factors. The use of communal items by both humans and animals, such as dog toys and computer terminals may pose a risk of infection and/or AMR transmission if thorough cleaning is not conducted between uses. Similarly, the high number of AMR genes found in the consulting table swabs suggest the potential for transmission, especially if they are used continually and not cleaned thoroughly in between uses. The non-slip rubber surface on these tables was observed to be covered in small scratches, presumably from animal claws, which would make them difficult to comprehensively clean. An alternative, more robust non-slip material could be utilised. The bin, kennel waste drain and sink plug hole samples also had high numbers of AMR genes identified. These data may suggest a call for more regular, or more thorough, deep cleaning of these areas, to reduce the risk of transmission to both humans and animals.

Hospital areas with fewer AMR genes identified were those with lower or no animal presence and/or movement, such as the pharmacy, imaging room and isolation ward, although as sampling was done at a single time point, the effect of before and after cleaning could not be elucidated. Another area of low animal movement and no AMR genes were recovered was the cat waiting area. In comparison, 22 AMR genes were identified in the sample from the dog waiting area. This may be due to the fact that cats will likely be in cat carriers whilst waiting, whereas dogs will be on the floor, or on the seats. With a large volume of pets waiting in these areas, cleaning cannot be undertaken between every patient, which strengthens the argument that high animal-movement areas, especially of dogs, are likely to have more AMR genes present in the environment, making them higher transmission risk areas.

The clinical pathogens most commonly identified within the environmental metagenomic samples were *E. faecalis, E. coli*, the Moraxella group*, S. pseudintermedius*, CoNS, *K. pneumoniae* and *P. aeruginosa*. These species can often harbour high numbers of resistance genes, as well as the ability to possess and transmit mobile genetic elements such as plasmids. Their presence in the veterinary hospital environment poses a potential risk for pathogen transmission, and the provision of a reservoir for AMR genes of importance, such as *mecA*, which may circulate between non-pathogenic and pathogenic species.

This study demonstrates that metagenomic sequencing analysis can be a useful tool for broad environmental surveillance of AMR genes and potentially pathogenic species, and may be impactful in terms of identifying IPC risk and areas for intervention. Whilst the cost may mean it is used in a targeted manner to guide practice rather than routinely, metagenomic surveillance for AMR in (human) clinical environments is gaining traction (69). There are limitations however, as lack of sensitivity/resolution makes it difficult to identify more detailed relationships, such as which species carry which plasmids and non-chromosomal AMR genes. This makes it difficult to identify whether it is the pathogenic, or environmental/commensal species, with MDRO profiles. Without specific enrichment or isolation of a species for whole genome sequencing, it is also not possible to identify transmission patterns of potential healthcare-associated infections, however, use of metagenomic sequencing can help to pinpoint areas of interest, and thus reduce potential sampling and costs for secondary, more detailed studies. Further considerations are the difficulties in ensuring standardization of sample collection and the ability to process some sample types, such as alcohol- and oil-based gels, as they are not compatible with DNA extraction kits.

## Conclusions

The results of this study should be placed in context; the environment, including that of a veterinary hospital, is not anticipated to be sterile and the identification of bacterial organisms including those with AMR genes, plasmids and MDRO phenotypes is not unexpected. As might be anticipated, our study suggested that high animal-handling and animal-movement areas, such as the preparation and consultation rooms, as well as areas in which dogs are present, are more likely to harbour AMR genes, plasmids and potentially pathogenic species. This information can be of clinical importance when considering infection prevention and control (IPC) measures however, and these findings would support the consideration of risk-based assessments to determine the frequency of cleaning and disinfection protocols within given areas. Surfaces that come into contact with animals or animal fluids such as consultation tables and bins should be considered intervention targets, together with the importance of measures such as hand hygiene to reduce the risk of direct and indirect transmission within the hospital environment.

## Abbreviations

AMR: Antimicrobial resistance

AST: Antibiotic susceptibility test

CoNS: Coagulase negative Staphylococci

CRE: Carbapenem resistant Enterobacteriales

ESBL: Extended-spectrum beta-lactamase

HAI: Healthcare associated infection

HHAI: Human-healthcare associated infection

LMIC: Low- and middle-income country

MDR: Multi-drug resistance

MDRO: Multi-drug-resistant organism

MIC: Minimum inhibitory concentration

MRS: Methicillin-resistant Staphylococci

MRSA: Methicillin-resistant *Staphylococcus aureus*

MRSP: Methicillin-resistant *Staphylococcus pseudintermedius*

ICR: Inducible clindamycin resistance

IPC: Infection prevention and control

OH: One Health

ONT: Oxford Nanopore Technologies

SD: Standard deviation

Spp: species

## Data availability statement

All data is Open Access and is available in the supplementary materials. All genetic fasta data obtained in this study is available on the European Nucleotide Archive, under the Project accession number PRJEB84924.

## Ethical approval and consent to participate

All clinical data were fully anonymised by the host veterinary practice and so patient/owner consent was not required.

## Author contributions

EQW, TDM, ADM and LE conceptualised the study, EQW, ADM and LE developed the environmental sampling protocol. ADM obtained clinical data, LE collected, extracted and sequenced metagenomic data. LE, SR, EQW, SMF and RJ analysed data. LE wrote the manuscript, all authors edited the manuscript.

