## supplementary data for "A metagenomic approach to One Health surveillance of antimicrobial resistance in a UK veterinary centre"

Supplementary materials

### 1. Veterinary Hospital floor plan

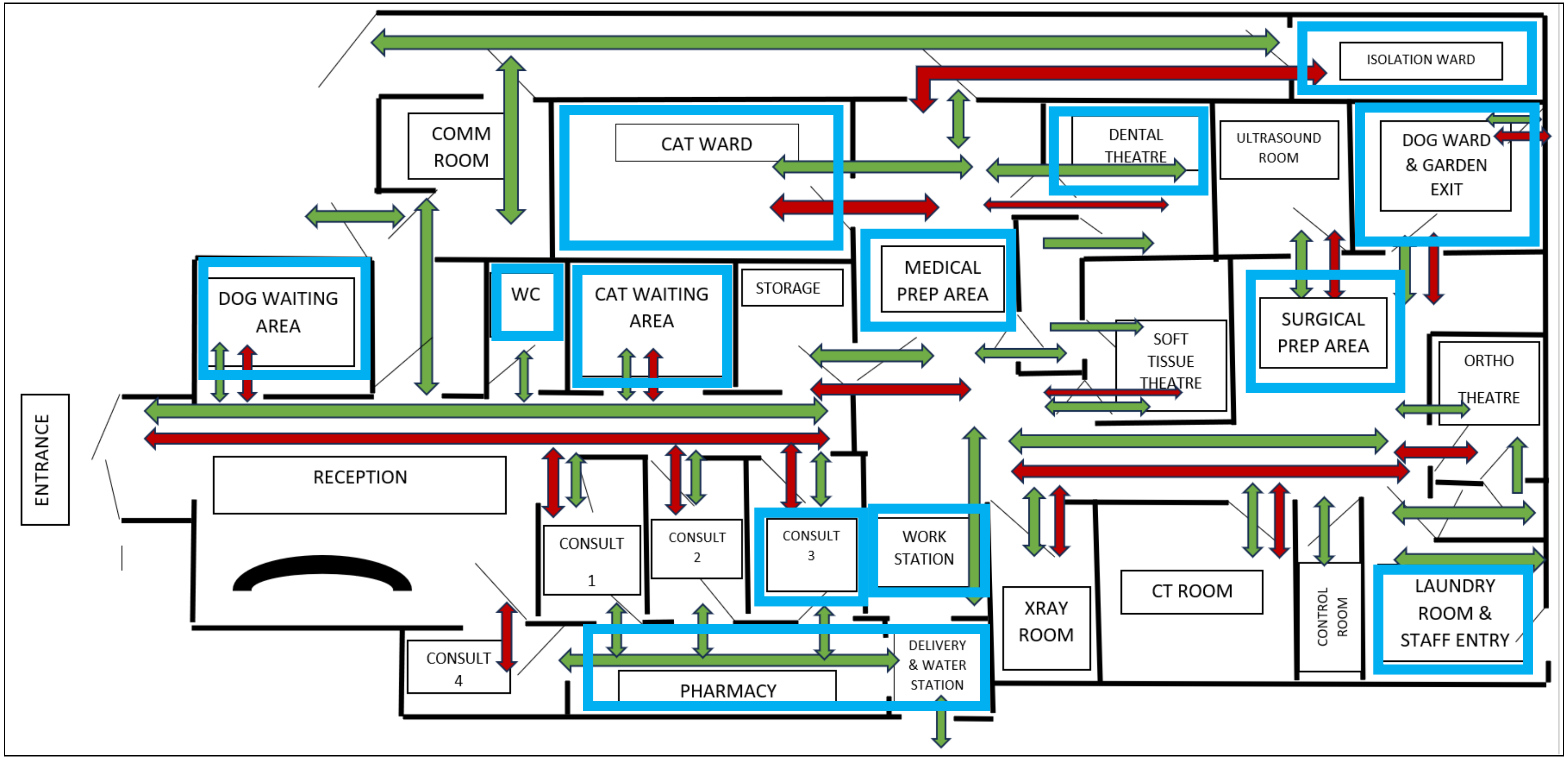


Layout of Small Animal Veterinary Hospital where samples were taken (not to scale). Green arrows represent human flow of movement. Red arrows show animal flow of movement. Areas highlighted by blue boxes were the rooms sampled.

#### 2. Sample data and accession numbers

BioProject number: PRJEB84924

| Sample number | Room | Description | Sample type | Total mapped reads | Fastq file size | Sample ID |
|---|---|---|---|---|---|---|
| 1 | Dog ward | Sink - plug hole | Stick swab | 0 | 0 KB | No data, not uploaded |
| 2 | Dog ward | Kennel waste drain | Wastewater | 1,349,483 | 961 MB | ERS22966141 |
| 3 | Dog ward | Communal toys | Sponge swab | 544,391 | 1.04 GB | ERS22966142 |
| 4 | Dog ward | Computer terminal | Sponge swab | 0 | 21.09 MB | ERS22966143 |
| 5 | Dog ward | Bins | Sponge swab | 981,862 | 698 MB | ERS22966144 |
| 6 | Dog ward | Consulting table | Sponge swab | 341,631 | 456 MB | ERS22966145 |
| 7 | Surgical preparation area | Sink - plug hole | Stick swab | 9,394,133 | 5.55 GB | ERS22966146 |
| 8 | Surgical preparation area | Consulting table | Sponge swab | 350,816 | 229 MB | ERS22966147 |
| 9 | Surgical preparation area | Computer terminal | Sponge swab | 52,939 | 239 MB | ERS22966148 |
| 10 | Surgical preparation area | Bins | Sponge swab | 2,399,821 | 2.66 GB | ERS22966149 |
| 11 | Laundry room | Sink - plug hole | Stick swab | 1,384,049 | 368 MB | ERS22966150 |
| 12 | Laundry room | Waste pipe and water catch tray | Wastewater | 1,116,071 | 598 MB | ERS22966151 |
| 13 | Imaging room | Ultrasound probe | Sponge swab | 5,974 | 6.18 MB | ERS22966152 |
| 14 | Dog ward | Door handles | Sponge swab | 31 | <1 KB | ERS22966153 |
| 15 | Back yard | Outside open water pipe | Wastewater | 115 | 20.53 KB | ERS22966154 |
| 16 | Medical preparation area | Sink - plug hole | Stick swab | 8,990,190 | 5.77 GB | ERS22966155 |
| 17 | Medical preparation area | Dog washing table | Sponge swab | 290 | 1.64 GB | ERS22966156 |
| 18 | Medical preparation area | Bins | Sponge swab | 2,944,186 | 2.62 GB | ERS22966157 |
| 19 | Medical preparation area | Consulting tables | Sponge swab | 10,991,957 | 4.9 GB | ERS22966158 |
| 20 | Medical preparation area | Computer terminal | Sponge swab | 28,521 | 12.92 MB | ERS22966159 |
| 21 | Medical preparation area | Intubation/oxygen tubes | Sponge swab | 3,200 | 17.11 MB | ERS22966160 |
| 22 | Medical preparation area | Thermometers | Sponge swab | 138,872 | 78.31 MB | ERS22966161 |
| 23 | Dental theatre | Table and equipment | Sponge swab | 11,592 | 60.12 MB | ERS22966162 |
| 24 | Dental theatre | Door handles | Sponge swab | 62 | 374 KB | ERS22966163 |
| 25 | Cat ward | Sink - plug hole | Stick swab | 654,306 | 493 MB | ERS22966164 |

| | | | | | | |
|---|---|---|---|---|---|---|
| 26 | Cat ward | Bins | Sponge swab | 367,562 | 1.32 GB | ERS22966165 |
| 27 | Cat ward | Consulting table | Sponge swab | 3,280 | 6.83 MB | ERS22966166 |
| 28 | Cat ward | Door handles | Sponge swab | 0 | 0 KB | No data, not uploaded |
| 29 | Cat ward | Computer terminal | Sponge swab | 0 | 0 KB | No data, not uploaded |
| 30 | Pharmacy/drinking room | Drinking fountain water tray | Wastewater | 2,027,394 | 3.17 GB | ERS22966167 |
| 31 | Veterinarian workstation room | Computer stations | Sponge swab | 2,176,452 | 845 MB | ERS22966168 |
| 32 | Veterinarian workstation room | Door handles | Sponge swab | 165,271 | 189 MB | ERS22966169 |
| 33 | Pharmacy/drinking room | Bins | Sponge swab | 14,369,377 | 3.44 GB | ERS22966170 |
| 34 | Pharmacy/drinking room | Water in mop bucket | Wastewater | 370,453 | 120 MB | ERS22966171 |
| 35 | Isolation ward | Door handles | Sponge swab | 41,283 | 22.92 MB | ERS22966172 |
| 36 | Reception toilet | Door handles | Sponge swab | 107,957 | 35.13 MB | ERS22966173 |
| 37 | Reception toilet | Toilet flush | Sponge swab | 6,611 | 3.42 MB | ERS22966174 |
| 38 | Reception toilet | Sink - plug hole | Stick swab | 190,649 | 264 MB | ERS22966175 |
| 39 | Cat reception area | Seating area | Sponge swab | 0 | 382 MB | ERS22966176 |
| 40 | Dog reception area | Seating area | Sponge swab | 56,017 | 136 MB | ERS22966177 |
| 41 | Consulting room 3 | Sink - plug hole | Stick swab | 55,255 | 23.45 MB | ERS22966178 |
| 42 | Consulting room 3 | Computer terminal | Sponge swab | 0 | 107 KB | ERS22966179 |
| 43 | Consulting room 3 | Door handles | Sponge swab | 29,839 | 16.58 KB | ERS22966180 |
| 44 | Consulting room 3 | Consultation tables | Sponge swab | 5,770,761 | 3.01 GB | ERS22966181 |
| 45 | Consulting room 3 | Bin | Sponge swab | 2,528,700 | 1.02 GB | ERS22966182 |
| 46+47 | Imaging room | Old (in use) ultrasound gel | Liquid sample | 0 | 11.97 KB | ERS22966183 |
| 48 | Imaging room | New (unopened) ultrasound gel #1 | Liquid sample | 0 | 43.73 KB | ERS22966184 |
| 49 | Imaging room | New (unopened) ultrasound gel #2 | Liquid sample | 0 | 0 KB | No data, not uploaded |
| 50 | Imaging room | Petroleum jelly | Liquid sample | 0 | 33.63 KB | ERS22966185 |

### 3. Swabbing procedures

| Sample type | Swab type | Size/volume | Process |
|---|---|---|---|
| Bins | Sponge swab | ~5 $cm^2$ | The lid and front of the bins were swabbed (sampling areas were selected at random) |
| Communal toys | Sponge swab | ~5 $cm^2$ | A sampling area was selected at random, from a toy that was also selected at random from the box |
| Computer terminal | Sponge swab | ~5 $cm^2$ | The mouse and randomly selected keys on the keyboard were swabbed |

| | | | |
|---|---|---|---|
| Consulting table | Sponge swab | ~5 cm$^2$ | A sampling area was selected at random from the surface of the table |
| Dog washing table | Sponge swab | ~5 cm$^2$ | A sampling area was selected at random from the surface of the table |
| Door handles | Sponge swab | Front and back of handle | The front and back of the door handle was swabbed |
| Drinking fountain water tray | Wastewater | 500 µL | The water in the tray was gently mixed to ensure an even suspension, then the sample was taken using a Pasteur pipette |
| Intubation/oxygen tube | Sponge swab | ~5 cm$^2$ | A sampling area was selected at random from the surface of a randomly selected intubation tube |
| Kennel waste drain | Sponge swab | 500 µL | The sample was taken using a Pasteur pipette, close to where the drain exited the room (so all kennels would have drained through the sample site) |
| Outside open water pipe drain | Gel sample | 500 µL | Wastewater was removed using a Pasteur pipette from the grate above the water line |
| Petroleum jelly | Wastewater | ~500 µL | A Pasteur pipette was used to remove the petroleum jelly from the container. Due to its viscous nature and poor water solubility (it did not mix with the DNA/RNA shield), the sample size was approximate |
| Seating area | Sponge swab | ~5 cm$^2$ | A sampling area was selected at random from the surface of the benches |
| Sink plug holes | Sponge swab | 500 µL | We did not have permission to remove the U-bends, so 500 µL wastewater was removed using a Pasteur pipette from the plug hole above |
| Table and equipment (dental theatre) | Sponge swab | ~5 cm$^2$ | A sampling area was selected at random from the surface of the table, as well as swabbing the switches that control the table and equipment |
| Thermometer | Sponge swab | Front and back of thermometer | The front and back of the thermometer was swabbed |
| Toilet flush | Sponge swab | Top and bottom of flush handle | The top and bottom of flush handle was swabbed |
| Ultrasound gel | Sponge swab | ~500 µL | A Pasteur pipette was used to remove the ultrasound gel from the container. The sample was then mixed with DNA/RNA Shield in a screw cap tube |
| Ultrasound probe | Wastewater | ~5 cm$^2$ of probe surface | The front, back and surface of the probe itself was swabbed |
| Waste pipe and water catch tray | Wastewater | 500 µL | Wastewater was removed using a Pasteur pipette from the waste pipe water catch tray |
| Water in mop bucket | Sponge swab | 500 µL | The water in the bucket was gently mixed to ensure an even suspension, then the sample was taken using a Pasteur pipette |

### 4. Commonly seen resistances for the clinical species seen in this study

| **Species** | **Commonly seen resistances** |
|---|---|
| Coagulase negative staph | Penicillin, oxacillin, gentamicin [40] |
| *Enterococcus faecalis* | Cephalosporins, lincosamides, clindamycin, aminoglycosides and some synthetic β-lactams [41] |
| Enterococcus spp | Cephalosporins, aminoglycosides, trimethoprim-sulfamethoxazole, Clindamycin, Lincosamides, Penicillinase-resistant penicillins, Bacitracin [41] |
| *Escherichia coli* | Tetracycline, sulfonamides, streptomycin, and ampicillin. *E. coli* found in animal hosts may also show resistances to phenicols, trimethoprim and Fosfomycin [42] |
| *Hafnia alvei* | Penicillin, oxacillin, amoxicillin plus clavulanic acid, colistin, narrow- and extended-spectrum cephalosporins [43], [44] |
| *Klebsiella oxytoca* | Fluoroquinolones, erythromycin, tetracycline, chloramphenicol, cefepime [45], [46] |
| *Klebsiella pneumoniae* | Ampicillin and some cephalosporins [47] |
| Moraxella group | vancomycin and trimethoprim. *M. catarrhalis* commonly produces a β lactamase [48], [49] |
| *Morganella morganii* | Oxacillin, Ampicillin, Amoxicillin, most first- and second-generation cephalosporins, Macrolides, Lincosamides, Glycopeptides, Fosfomycin, Fusidic acid, Colistin [50], [51] |
| *Pasteurella multocida* | *P. multocida* is generally thought to be susceptible to the majority of antimicrobials [52] |
| *Pleuralibacter gergoviae* | Penicillins, macrolides, lincosamides, streptogramins, rifampicin, fusidic acid, Fosfomycin, cefoxitin (likely due to β-lactamase production) [53], [54] |
| *Proteus mirabilis* | Tetracycline, nitrofurantoin, polymyxin, colistin, reduced susceptibility to imipenem [55] |
| *Pseudomonas aeruginosa* | β-lactam and penem antibiotics [56] |
| Salmonella spp | Resistance to antimicrobials varies between *Salmonella* serotypes [57] |
| *Serratia fonticola* | Amoxicillin, ticarcillin [58] |
| *Staphylococcus aureus* | Methicillin-sensitive Staphylococcus aureus (MSSA) or methicillin-resistant Staphylococcus aureus (MRSA), which is resistant to penicillin, methicillin, aminoglycosides [59] |
| *Staphylococcus haemolyticus* | Noted to be one of the most resistant CoNS. Penicillins, cephalosporins, macrolides, quinolones, tetracyclines, aminoglycosides, glycopeptides, fosfomycin [60], [61] |
| *Staphylococcus lentus* | Macrolides, lincosamides, streptogramins, tetracycline [62], [63] |
| *Staphylococcus pseudintermedius* | Penicillin, tetracycline, erythromycin, amikacin, gentamicin, enrofloxacin [64], [65] |
| *Staphylococcus saprophyticus* | Novobiocin [66] |
| *Staphylococcus schleiferi* | Penicillin, fluoroquinolones, methicillin [67] |

| | |
|---|---|
| *Streptococcus canis* | Some strains have shown resistance to vancomycin, linezolid. Further strains may also show resistance to β-lactams [68], [69] |
